# Partial absence of PD-1 expression by tumor-specific CD8^+^ T cells in EBV-driven lymphoepithelioma-like carcinoma: a case report

**DOI:** 10.1101/696278

**Authors:** Yannick Simoni, Etienne Becht, Shamin Li, Chiew Yee Loh, Joe Poh Sheng Yeong, Tony Kiat Hon Lim, Angela Takano, Daniel S.W. Tan, Evan W. Newell

## Abstract

Lymphoepithelioma-like carcinoma (LELC) is an uncommon lung cancer, typically observed in young, non-smoking Asian populations. LELC is associated with Epstein-Barr virus (EBV) infection of lung tumor cells of epithelial origin, suggesting a carcinogenic role of EBV as observed in nasopharyngeal carcinoma (NPC). Here, we studied the antigen specificity and phenotype of CD8^+^ tumor infiltrating lymphocytes (TILs) in one LELC patient positive for EBV infection in lung tumor cells. Using MHC class I tetramers, we detected two populations of EBV-specific CD8^+^ TILs, which can be considered as tumor-specific CD8^+^ T cells, in the tumor of this patient. Transcriptomic analyses of these two populations reveal their distinct exhausted profiles and polyclonal TCR repertoire. High dimensional analyses at single cell level using mass cytometry showed showed that populations of tumor specific CD8^+^ TILs are phenotypically heterogeneous, although they consistently express CD39. Unexpectedly, although the LELC tumor cells expressed abundant PD-L1, these tumor-specific CD8^+^ TILs mostly did not express PD-1, suggesting that anti-PD1/PD-L1 immunotherapy may not be an appropriate strategy for disinhibiting EBV-specific cells for the treatment of LELC patients. These results might also help to explain low rates of checkpoint blockade immunotherapy response for NPC, despite the antigenicity of EBV for both tumor types.

## Introduction

A better characterization of tumor-specific T cells remains an important challenge to improve efficacy of immune-based therapy [1]. However, due to the presence of cancer unrelated “bystander” T cells in the tumor [2, 3] and the difficulty to identify tumor-specific T cells (e.g. neoantigen prediction) [4], study of these cells remains challenging in cancer. Human cancers associated with viral infections represent approximatively 10% of the worldwide cancer incidence [5]. Due to the presentation by MHC class I molecule of well characterized viral antigens by tumor cells, virus-associated cancers are relevant to study tumor-specific T cells by studying virus-specific T cells in the tumors.

Oncogenic virus such as hepatitis B virus (HBV) in liver cancer, human papillomavirus (HPV) in cervical cancer or Epstein-Barr virus (EBV) in B cells lymphoma, disturb biological pathways to replicate in tumor infected cells and escape the immune surveillance [6, 7]. Importantly, none of these viral infections on their own is sufficient to induce cancer but is closely related to genetic susceptibility (e.g. single nucleotide variant, driver mutation) and environmental factors (e.g. viral co-infection, dietary) [7–9]. Lymphoepithelioma-like carcinoma (LELC) is a rare cancer, characterized by a massive infiltration of lymphocytes [10]. The term LELC refers to its histological resemblance with the lymphoepithelioma tumor observed in Nasopharyngeal carcinoma (NPC). The first case of LELC in lung was reported in 1987 [11] and has been observed in several other tissues [12] such as gastrointestinal tract [13], salivary glands [14], skin [15], breast [16] or vagina [17]. Because of the absence of driver mutation (i.e. EGFR, ALK) in the majority of patients with LELC of the lung [18], effective treatment options are limited (e.g. TKI drugs). A strong association between LELC in lung and EBV infection has been observed in Asian populations [19]. Based on observations made for NPC, the mechanistic model suggests that EBV virus is not an initiating factor in the oncogenic process, but a tumor-promoting agent. Loss of heterozygosity in epithelial cells associated with genetic (e.g. Asian ethnicity) and environmental factors (e.g. salt fish) is an early event in the pathology of the disease. Infection of these DNA-damaged epithelial cells by EBV, followed by expression of EBV latent gene, will provide growth and survival advantages to these cells (e.g. BCL2 overexpression) and lead to the development of carcinoma that may finally result in metastasis [7, 9]. Study of EBV-specific T cells in EBV-associated tumors (i.e. NPC, LELC) represent an important step to develop more efficient therapeutic tools to treat these cancers, such as TCR engineered T-cells targeting EBV epitopes presented by tumor cells, or improved checkpoint blockade immunotherapy (e.g. anti-PD-1, anti-PD-L1).

Here, using MHC class I tetramers to identify EBV-specific CD8^+^ TILs in a patient with LELC, we reported the simultaneous detection of two EBV-specific populations. Although these two populations shared common characteristics, we highlight the polyclonality of their T cell receptors, their heterogeneous exhausted/dysfunctional profile, and the unexpected partial absence of PD-1 surface protein expression.

## Results

### Identification of EBV-specific CD8^+^ TILs in EBV driven LELC cancer

A 69 year old Asian woman (patient A311), non-smoker, was diagnosed with a lung cancer. Histological examination from resected tumor of this patient indicated a lymphoepithelioma-like carcinoma (LELC) structure with an abundant infiltration of immune cells. Because LELC of the lung is strongly associated with the presence of Epstein-Barr virus (EBV) in tumor cells [19], we performed EBV-encoded RNA in-situ hybridization (EBERish) staining on tumor tissue sections. We confirmed the presence of EBV viral RNA in the tumor cells of this patient (Figure 1A). Since EBV proteins are presented by MHC class I molecules from the tumor cells, CD8^+^ TILs specific for EBV epitopes can be considered as tumor specific. Based on the HLA-A alleles expressed by this patient, we screened for EBV-specific CD8^+^ TILs using HLA-A*24:02 tetramers loaded with different EBV epitopes (see methods). We detected a high frequency of two EBV-specific CD8^+^ TIL populations recognizing the BMFL1 (DYNFVKQLF – 0.70%) and BRFL1 (TYPVLEEMF – 0.43%) epitopes (Figure 1B). These two proteins are implicated in the early lytic cycle of EBV [7], strongly suggesting that EBV is producing virions in tumor cells. We did not detect any cells specific for EBV latency cycle associated antigens in this patient despite testing for epitopes derived from three latency associated gene products (LMP2, EBNA3A or EBNA3B, see discussion).

**Figure 1.**
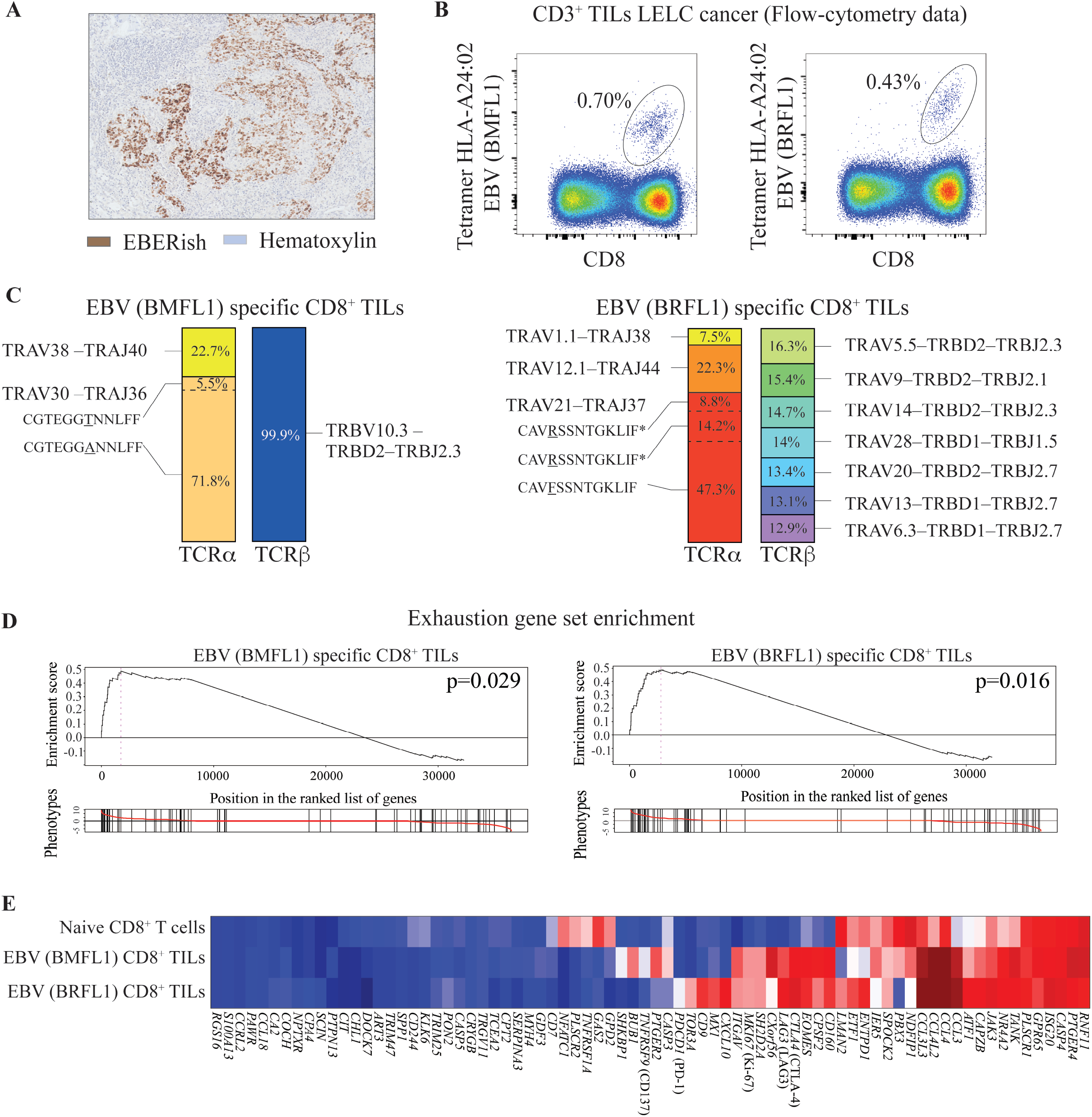
Identification of EBV-specific CD8^+^ TILs in EBV infected LELC of the lung. **A**. Immunohistochemistry staining of the lung from the lymphoepithelioma-like carcinoma (LELC) patient stained with hematoxylin (blue) and Epstein-Barr virus-encoded small RNA in-situ hybridation (EBERish – brown). **B**. Flow-cytometry dot plot representing two cell populations specific for EBV-derived peptides (parts of the BRFL1 and BMFL1 proteins) infiltrating LELC tumor from patient A311. Frequency of tetramer-positive cells among CD3^+^ TILs. **C**. TCRα and TCRβ clones repertoire of EBV(BMFL1) and EBV (BRFL1) specific CD8^+^ TILs from LELC lung cancer (patient A311). * Same TCRα amino acid sequence but different TCRα nucleotide sequence. **D**. Enrichment of the gene set for exhausted T cells in EBV(BMFL1) or EBV (BRFL1) specific CD8^+^ TILs. Gene position on the left indicates enrichment in EBV (BMFL1) or EBV (BRFL1) specific CD8^+^ TILs. Gene position on the right indicates enrichment in naive CD8^+^ T cells. **E**. Heatmap of gene set for exhaustion shown in D.

### EBV-specific CD8^+^ TILs in EBV-driven LELC have a polyclonal TCR repertoire and are exhausted

We FACS sorted and analyzed the transcriptomic profile of both EBV-specific CD8^+^ TIL populations in this patient. These cells were analyzed without prior expansion that could alter the TCR repertoire and transcriptomic profile. We first studied the TCR repertoire of each population. Both EBV-specific CD8^+^ TIL populations were polyclonal for their TCRα, with three and four different TCRα clones detected for each epitope (Figure 1C). Interestingly, BMFL1 tetramer^+^ cells expressed only one TCRβ clone. In contrast, for BRFL1 tetramer^+^ cells, seven different TCRβ clones were detected at similar frequency (Figure 1C). This result highlights the TCR polyclonality of CD8^+^ TILs specific for each individual epitope and suggest that a skewed TCRβ repertoire observed in tumors do not necessarily translate an expansion of tumor-specific CD8^+^ TILs, as observed for BRFL1 tetramer^+^ cells [20]. Furthermore, polyclonal TCRα/β observed raises questions about the best strategy (i.e. TCRα/β choice) to design TCR engineered T-cells therapy (see Discussion).

Gene set enrichment analysis revealed a significant exhausted profile of both EBV-specific CD8^+^ TIL populations (Figure 1D). Although both populations similarly expressed genes involved in dysfunction/exhaustion status (i.e. *CTLA4, LAG3, ENTPD1*) or proliferation (i.e. *MKI67*), we surprisingly observed a dichotomy for some other genes. For example, the inhibitory receptor *PDCD1* (PD-1) was only detected in EBV(BRFL1)-specific CD8^+^ TILs at low level. Despite being in the same tumor microenvironment and being specific for the same virus, the two populations showed a distinct exhaustion profile at the transcriptome level (see Discussion). To confirm these differences, we analyzed surface markers expressed by both populations at the single cell level using mass-cytometry.

### EBV-specific CD8^+^ TILs in EBV-driven LELC are phenotypically heterogenous and express CD39

We developed a mass-cytometry panel of metal-labeled antibodies to analyze TILs from resected lungs (see methods). This panel included a broad range of phenotypic markers that allow to simultaneously measure the expression of markers associated with T cell differentiation, activation, tissue residence, and the dysfunctional/exhausted status of each cell (Figure 2A). For high-dimensional assessment of tetramer^+^ CD8^+^ TILs heterogeneity, we used Uniform Manifold Approximation and Projection (UMAP), which accounts for non-linear relationships between markers and projects high dimensional data into two dimensions (called UMAP1 and UMAP2) by making a pair-wise comparison of cellular phenotypes to optimally plot similar cells nearby to each other [21, 22]. In the tumor microenvironment, the presence of different clusters on the UMAP map for each of the EBV-specific CD8^+^ TIL populations highlight their phenotypic heterogeneity (see Clusters 1, 2, 3 and 4 – Figure 2B). Even being phenotypically heterogeneous, both EBV-specific CD8^+^ TIL populations shared several similar characteristics. Both displayed an effector phenotype (CCR7^−^ CD45RO^+^) and expressed tissues resident memory markers (CD69, CD49a). Furthermore, as we previously reported, all cells expressed the marker CD39 [2]. The heterogeneity of both EBV-specific CD8^+^ TILs was driven by differential expression of integrin CD103 (cluster 4), the marker for senescence CD57 (cluster 1 and 2), and the adhesion molecule CD56 (cluster 2). Altogether, these observations show the phenotypic heterogeneity of two EBV-specific CD8^+^ TIL populations (see discussion).

**Figure 2.**
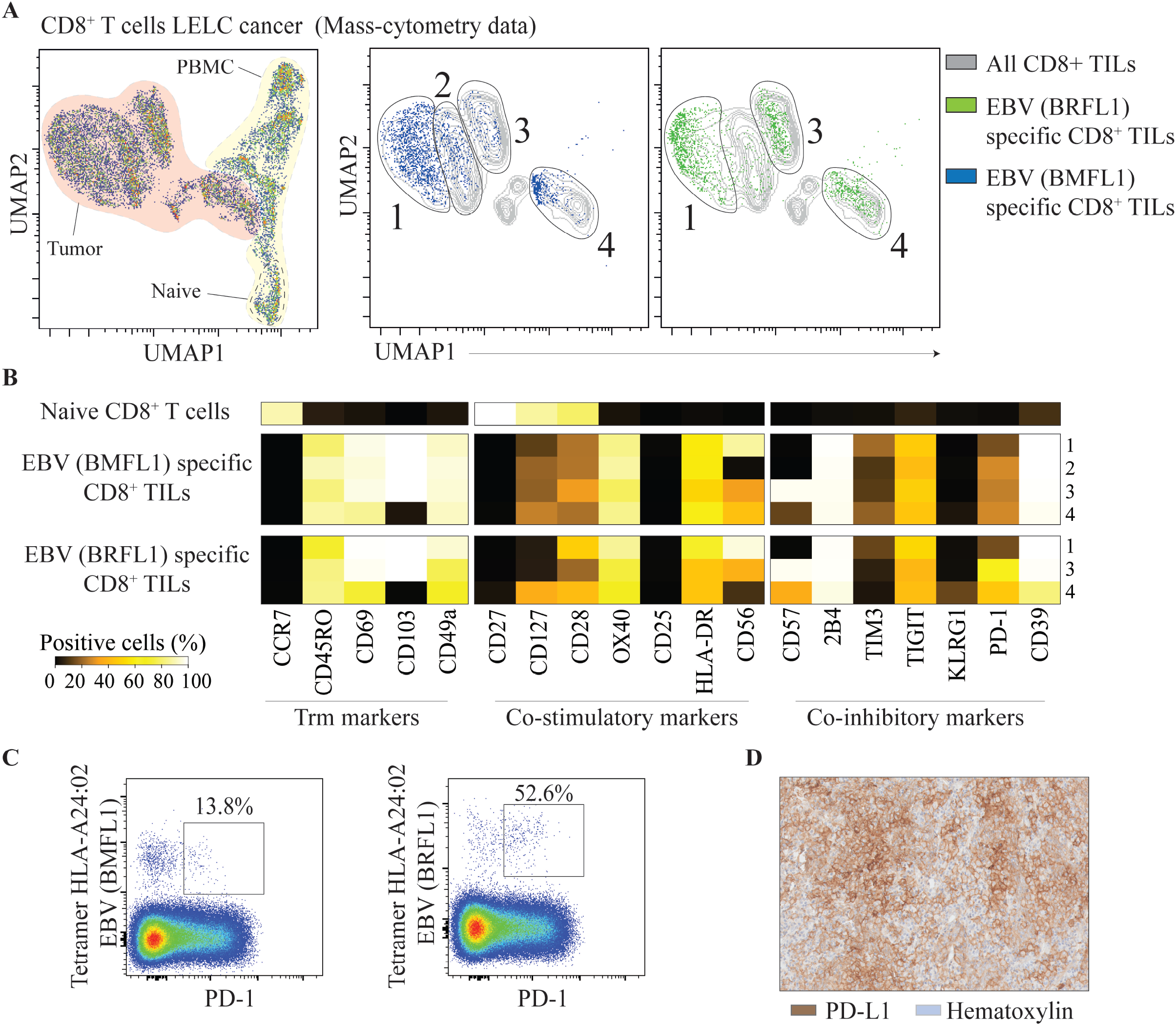
Tumor-specific CD8^+^ TILs in LELC are heterogenous and partially lack expression of PD-1. **A**. Uniform Manifold Approximation and Projection (UMAP) map of CD8+ T cells from blood (yellow) and tumor (red) isolated from patient A311. UMAP accounts for non-linear relationships between markers and projects high dimensional data into a low dimensional space (called UMAP1 and UMAP2) by making a pair-wise comparison of cellular phenotypes to optimally plot similar cells nearby each other. **B**. UMAP map of EBV(BMFL1–blue) or EBV (BRFL1–green) specific CD8+ T cells in blood and tumor. For each epitope, clusters (cells populations) were identified by manual gating. **C**. Based on the clusters identified in (B), frequencies of cells positive for each marker were calculated and represented as a heatmap **D**. Flow-cytometry dot plot representing expression of PD-1 by the two populations specific for EBV-derived peptides (parts of the BRFL1 and BMFL1 proteins) infiltrating LELC tumor from patient A311. Frequency of PD-1 positive cells among tetramer^+^ CD8^+^ TILs. **E**. Immunohistochemistry staining of the lung from the lymphoepithelioma-like carcinoma (LELC) patient stained with hematoxylin (blue) and PD-L1 (brown).

### Expression of PD-1 is partially absent on EBV-specific CD8^+^ TILs from the LELC patient

Expression of the inhibitory receptor PD-1 is known to inhibit tumor-specific CD8^+^ T cells activation [23, 24]. Treatment of cancer patients with anti-PD-1 or anti-PD-L1 has succesfully induced tumor regression in some cases. However, anti-PD-1 efficacy is variable across patients and cancer types [25]. Recently, we showed that PD-1 can also be expressed by bystander T cells in the tumors [2], making this receptor not exclusively expressed by tumor-specific cells, and raising questions about the effect of anti-PD-1 treatment on these cells. Surprisingly, the two EBV-specific CD8^+^ TILs infiltrating this EBV-driven LELC tumor only partially expressed PD-1, 13.8% and 52.6% of positive cells respectively for BMFL1 and BRFL1 epitopes (Figure 2D). Furthermore, the PD-1 intensity of these cells is weak compared to PD-1 expression by tumor associated antigen or neoantigen-specific CD8 TILs (Supplementary Figure 1). Nonetheless, we noted that the tumor cells expressed high level of PD-L1 in this tumor (Figure 2E). Because of the weak and partial PD-1 expression on EBV-specific CD8^+^ TILs infiltrating EBV-driven LELC, our data suggest that anti-PD-1 treatment for EBV-driven LELC may not be able to disinhibit these EBV and tumor-specific cells.

### Discussion

In this study, we report the identification of two EBV-specific CD8^+^ T cell populations infiltrating an EBV-driven LELC of the lung. Tumor specific CD8^+^ cells have been characterized by an exhausted/dysfunctional signature, which can be reverted by immunotherapy [26]. However, CD8^+^ TILs are highly heterogenous across tumors, patients and cancer types [2, 27]. This heterogeneity across patients can be explained by tumor cell-intrinsic factors shaping the tumor immune microenvironment and finally influencing the outcome of immunotherapy [28]. Recently, we demonstrated that intra-tumor heterogeneity can be in part explained by the presence of tumor-specific (exhausted phenotype, CD39^+^) and bystander CD8^+^ TILs [2]. Here, in the same tumor-microenvironment, we report distinct exhaustion signatures between two tumor-specific CD8^+^ TIL populations. Furthermore, each of these two tumor-specific populations is phenotypically distinct at the single-cell level. We previously reported similar observations in a mouse tumor model [29]. This heterogeneity in the same tumor environment suggests that unidentified factors (e.g. tumor-epitope availability, spatial localization or TCR repertoire diversity/uniformity) influence the acquisition of different exhausted/dysfunctional profiles in the same tumor microenvironment. As reported previously, LELC tumor cells express high level of PD-L1 [30], making LELC cancer a potential good candidate for anti-PD-1 or anti-PD-L1 immunotherapy. However, case report studies show variable effects of this treatment in LELC, one showing a partial response [31] and another report of a non-responder patient [32]. Surprisingly, we observed in our LELC patient that both tumor-specific CD8^+^ TILs only partially expressed PD-1 (13.8% and 52.6%). This observation could explain the partial absence of response of LELC patients to anti-PD-1 treatment. In this line, this observation strongly suggests that targeting PD-1–PD-L1 pathway by immunotherapy in LELC might not be the most appropriate strategy. Of note, relatively low rates of response to to anti-PD1 immunotherapy are being observed for NPC, which is also an EBV-related carcinoma [33][34]. We think it is interesting that EBV-related tumors do not show high response rates to checkpoint blockade immunotherapy despite the strong T cell immunogenicity of EBV. Our data showing that tumor infiltrating EBV-specific cells can display hallmarks of chronic antigen-stimulation and exhaustion even without PD-1 expression may help to explain this.

Effective treatment options in LELC are limited. The oncogenic role of EBV has been attributed to the expression of latent genes providing growth and survival benefit to DNA-damaged epithelial cells. Several therapeutic strategies have been developed to induce a cytotoxic CD8^+^ T cells (CTL) response targeting specifically the latent protein, such as autologous CTL infusion [35], vaccination with EBV-latent vector [36] or pulsed-dendritic cells [37, 38]. However, these strategies show moderate efficacy, with toxicity in some cases [39]. In our study, we did not detect CD8^+^ TILs specific for EBV latent protein among the three epitopes tested (LMP1, EBNA3A, EBNA3B), we only detected EBV-specific CD8^+^ TILs for lytic proteins (BMFL1 and BRFL1). Recently, transcriptomic analysis has highlighted that lytic genes can also be expressed in EBV-infected tumor cells [40]. Taken together, our observation suggests that targeting lytic proteins, in association with latent proteins, could be more efficient for autologous CTL infusion or vaccination strategies. In the last few years, chimeric antigen receptor (CAR) T cells therapy has shown promising efficacy in the treatment of B lymphoma [41, 42]. Similarly, TCR engineered T-cells targeting tumor epitopes have been developed and are used in phase I trials (i.e. MAGE3, NY-ESO) [43]. Engineered T-cells expressing a TCR targeting the latent EBV protein LMP1 presented by HLA-A*02:01 have been generated in a pre-clinical model [44]. Based on our observations, targeting lytic EBV epitopes presented by tumor cells could represent an interesting alternative strategy in EBV-driven cancer, due to the presence of the same EBV epitopes in different patients and the absence of these epitopes in the non-infected normal cells. However, our data highlight the polyclonal TCR repertoire of EBV-specific CD8^+^ TILs as well. Thus, more studies are needed to: 1) evaluate the efficiency of Engineered T-cells targeting lytic vs. latency associated EBV proteins, 2) evaluate the efficiency of monoclonal versus polyclonal TCR engineered T-cells. Taken together, our investigation of TIL specificities in the context of a virus-associated cancer should help inform the design and targets of future immunotherapeutic modalities

## Author contributions

Y.S. designed and did research, analyzed data and wrote the paper. S. L. Helped to analyze data and reviewing the paper. E.B. helped to analyze transcriptomic data. C.Y.L., helped to process samples. J.Y, A.T, T.K.H.L, D.S.W.T, provided samples and discussed data. E.W.N. initiated and led the project, developed scripts for CyTOF analysis and wrote the paper.

## Acknowledgments

The authors thank all members of E.N lab, Duan Kaibo and the bioinformatics group, Josephine Lum and the SIgN community, the Flow cytometry platform and the clinical research coordinators from NCCS. This study was funded by A-STAR/SIgN core funding and A-STAR/SIgN immunomonitoring platform funding.

## Experimental Procedures

### Human samples, Immunohistochemistry and cells isolation

PBMC and tumor samples were obtained from a 69 years old Asian woman (patient A311), non-smoker, with a lung cancer. Immunohistochemistry for H&E, PD-L1 and EBERish were performed on PFA fixed tissue sections [45]. Tumor single cell suspensions were prepared as previously described [46]. Briefly, tissues were mechanically dissociated in small pieces and incubated at 37ºC for 15 min DMEM + Collagenase IV (1 mg/ml) + DNAse (15 µg/ml). Digestion was stopped by addition of RPMI 5% FBS. Dissociated tissues were filtered and washed in RPMI 5% + DNAse (15 µg/ml) FBS. Single cell suspensions were cryopreserved in 90% FBS + 10% DMSO and stored in liquid nitrogen. The use of human tissues was approved by the appropriate IRBs, A*STAR, the Singapore Immunology Network.

### EBV-specific MHC class I tetramer and staining

HLA-A*24:02 monomer with a U.V cleavable peptide were produced as described previously [2]. Peptide exchange was performed using the following peptides specific of EBV virus: LMP2 (IYVLVMLVL), EBNA3 (RYSIFFDYM), BRLF1 (TYPVLEEMF), BMFL1 (DYNFVKQLF), EBNA3B (TYSAGIVQI). Each MHC class I monomer was tetramerized using streptavidin at the ratio 4:1 as described previously [2]. Frozen samples were thawed and washed. Cells were incubated for 1 hour at room temperature with the tetramer. Cells were then incubated with antibodies cocktail for 15mn and acquired on flow-cytometry.

### Mass-cytometry experiment

Purified antibodies lacking carrier proteins were conjugated to a heavy metal according to the protocol provided by Fluidigm Inc. Streptavidin was heavy metal-labeled as previously described [2]. Frozen samples were thawed and washed in RPMI 10% FBS + DNAse (15 μg/ml). Cells were incubated for 1 hour at room temperature with the tetramer cocktail. Then, cells were stained with Cisplatin (viability marker) at 5 µM in PBS for 5 min. Cells were then incubated with antibodies cocktail for 15mn and fixed in PFA 2% prior to CyTOF acquisition.

### mRNA sequencing data analysis

The paired-end RNA-seq reads from HiSeq 4000 were mapped to Human GRCh38/HG19 reference genome using STAR software tool. The mapped paired-end reads were summarized to gene-level using featureCounts v1.5.0-p1 software tool and with GENCODE v26 gene annotation. Genes with read count less than 5 in less than 2 samples in all cell populations were filtered out from further analysis. Limma-voom pipeline was used for differentially expressed gene (DEG) analysis. DEGs from comparisons between different cell populations were selected with Benjamini-Hochberg adjusted p-value of < 0.05. All analyses were done in R-3.1.2. We used the *HTSanalyzeR* package (v 2.26.0) to run GSEA on gene collections from the on the T cell exhaustion gene set [47, 48] filtered for gene sets with at least 20 genes present in our dataset. For GSEA we used 1000 permutations to estimate p-values and applied corrections for multiple tests using the Benjamini-Hochberg procedure. To measure TCR diversity, we converted the measured counts for each TCRα and TCRβ to frequencies.

### Data analysis and UMAP

After CyTOF acquisition, which was performed as previously described [29], any zero values were randomized using a uniform distribution of values between zero and minus-one using a R script. Note also that all other integer values measured by the mass cytometer are randomized in a similar fashion by default. The signal of each parameter was then normalized based on the EQ beads (Fluidigm) as previously described [49]. Samples were then used for UMAP analysis similar to that previously described [21]. In R, all data were transformed using the “logicleTransform” function (“flowCore” package) using parameters: w=0.25, t=16409, m=4.5, a=0 to roughly match scaling historically used in FlowJo.

---

**Supplementary Figure 1.**
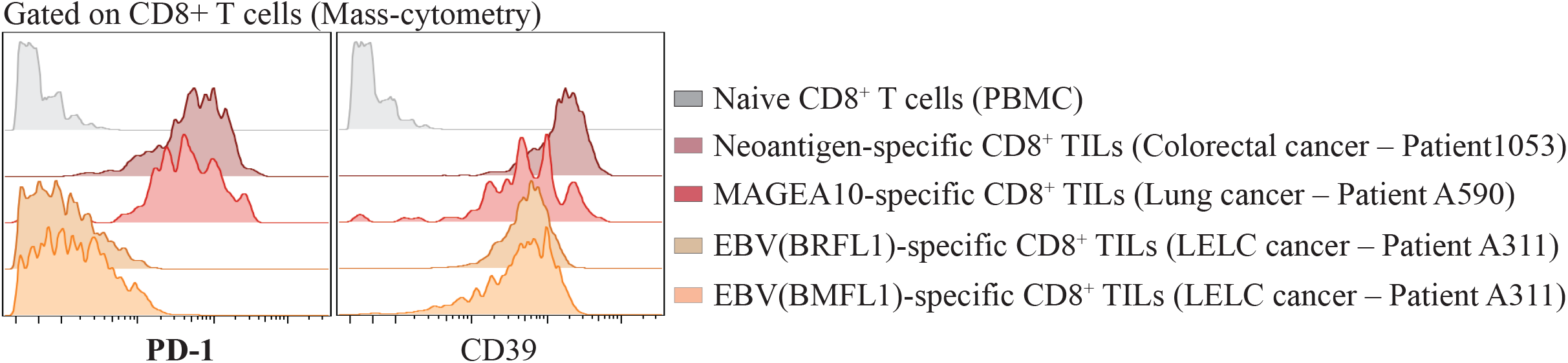
Expression of PD-1 by EBV, tumor associated antigen and neoantigen-specific CD8^+^ TILs in human cancer. Histogram representing PD-1 expression by tumor-specific CD8^+^ TILs identified by MHC class I tetramer. Naive CD8^+^ T cells (CCR7^+^ CD45RO^−^), Neoantigen-specific CD8^+^ TILs were identified using HLA-A*11:01 tetramer for mutAHR (GISQELPYK) (See Simoni Y, et al, Nature 2018), tumor associated antigens MAGEA10-specific CD8^+^ TILs were identified using HLA-A*02:01 tetramer (GLYDGMEHL).

## References

1. Newell, E.W. and E. Becht, High-Dimensional Profiling of Tumor-Specific Immune Responses: Asking T Cells about What They “See” in Cancer. Cancer Immunol Res, 2018. 6(1): p. 2–9.

2. Simoni, Y., et al., Bystander CD8(+) T cells are abundant and phenotypically distinct in human tumour infiltrates. Nature, 2018. 557(7706): p. 575–579.

3. Becht, E., E. W. Newell, and Y. Simoni, [Identification of tumor-specific CD8 T cells with a surface marker]. Med Sci (Paris), 2018. 34(12): p. 1032–1034.

4. Scheper, W., et al., Low and variable tumor reactivity of the intratumoral TCR repertoire in human cancers. Nat Med, 2019. 25(1): p. 89–94.

5. zur Hausen, H., Viruses in human cancers. Science, 1991. 254(5035): p. 1167–73.

6. El-Sharkawy, A., L. Al Zaidan, and A. Malki, Epstein-Barr Virus-Associated Malignancies: Roles of Viral Oncoproteins in Carcinogenesis. Front Oncol, 2018. 8: p. 265.

7. Young, L.S., L.F. Yap, and P.G. Murray, Epstein-Barr virus: more than 50 years old and still providing surprises. Nat Rev Cancer, 2016. 16(12): p. 789–802.

8. Wu, S., et al., Evaluating intrinsic and non-intrinsic cancer risk factors. Nat Commun, 2018. 9(1): p. 3490.

9. Shah, K.M. and L.S. Young, Epstein-Barr virus and carcinogenesis: beyond Burkitt’s lymphoma. Clin Microbiol Infect, 2009. 15(11): p. 982–8.

10. Ho, J.C., M.P. Wong, and W.K. Lam, Lymphoepithelioma-like carcinoma of the lung. Respirology, 2006. 11(5): p. 539–45.

11. Begin, L.R., et al., Epstein-Barr virus related lymphoepithelioma-like carcinoma of lung. J Surg Oncol, 1987. 36(4): p. 280–3.

12. Aurilio, G., et al., A possible connective tissue primary lymphoepithelioma-like carcinoma (LELC). Ecancermedicalscience, 2010. 4: p. 197.

13. Chen, P.C., et al., Epstein-Barr virus-associated lymphoepithelioma-like carcinoma of the esophagus. Hum Pathol, 2003. 34(4): p. 407–11.

14. Sun, X.N., et al., Lymphoepithelioma-like carcinoma of the submandibular salivary gland: a case report. Chin Med J (Engl), 2006. 119(15): p. 1315–7.

15. Cavalieri, S., et al., Lymphoepithelioma-like carcinoma of the skin. Int J Immunopathol Pharmacol, 2007. 20(4): p. 851–4.

16. Dinniwell, R., et al., Lymphoepithelioma-like carcinoma of the breast: a diagnostic and therapeutic challenge. Curr Oncol, 2012. 19(3): p. e177–83.

17. McCluggage, W.G., Lymphoepithelioma-like carcinoma of the vagina. J Clin Pathol, 2001. 54(12): p. 964–5.

18. Liang, Y., et al., Primary pulmonary lymphoepithelioma-like carcinoma: fifty-two patients with long-term follow-up. Cancer, 2012. 118(19): p. 4748–58.

19. Tay, C.K., et al., Primary pulmonary lymphoepithelioma-like carcinoma in Singapore. Ann Thorac Med, 2018. 13(1): p. 30–35.

20. Pandolfi, F., et al., Skewed T-cell receptor repertoire: more than a marker of malignancy, a tool to dissect the immunopathology of inflammatory diseases. J Biol Regul Homeost Agents, 2011. 25(2): p. 153–61.

21. Becht, E., et al., Dimensionality reduction for visualizing single-cell data using UMAP. Nat Biotechnol, 2018.

22. McInnes, L.H., J.; Melville, J.;, Umap: Uniform manifold approximation and projection for dimension reduction. 2018.

23. Nishino, M., et al., Monitoring immune-checkpoint blockade: response evaluation and biomarker development. Nat Rev Clin Oncol, 2017. 14(11): p. 655–668.

24. Ribas, A. and J.D. Wolchok, Cancer immunotherapy using checkpoint blockade. Science, 2018. 359(6382): p. 1350–1355.

25. Seidel, J.A., A. Otsuka, and K. Kabashima, Anti-PD-1 and Anti-CTLA-4 Therapies in Cancer: Mechanisms of Action, Efficacy, and Limitations. Front Oncol, 2018. 8: p. 86.

26. Wherry, E.J., T cell exhaustion. Nat Immunol, 2011. 12(6): p. 492–9.

27. Hashimoto, M., et al., CD8 T Cell Exhaustion in Chronic Infection and Cancer: Opportunities for Interventions. Annu Rev Med, 2018. 69: p. 301–318.

28. Li, J., et al., Tumor Cell-Intrinsic Factors Underlie Heterogeneity of Immune Cell Infiltration and Response to Immunotherapy. Immunity, 2018. 49(1): p. 178–193 e7.

29. Fehlings, M., et al., Checkpoint blockade immunotherapy reshapes the high-dimensional phenotypic heterogeneity of murine intratumoural neoantigen-specific CD8(+) T cells. Nat Commun, 2017. 8(1): p. 562.

30. Chang, Y.L., et al., PD-L1 is highly expressed in lung lymphoepithelioma-like carcinoma: A potential rationale for immunotherapy. Lung Cancer, 2015. 88(3): p. 254–9.

31. Kumar, V., et al., Response of advanced stage recurrent lymphoepithelioma-like carcinoma to nivolumab. Immunotherapy, 2017. 9(12): p. 955–961.

32. Kim, C., et al., Metastatic lymphoepithelioma-like carcinoma of the lung treated with nivolumab: a case report and focused review of literature. Transl Lung Cancer Res, 2016. 5(6): p. 720–726.

33. Ma, B.B.Y., et al., Antitumor Activity of Nivolumab in Recurrent and Metastatic Nasopharyngeal Carcinoma: An International, Multicenter Study of the Mayo Clinic Phase 2 Consortium (NCI-9742). J Clin Oncol, 2018. 36(14): p. 1412–1418.

34. Jain, A., W.K. Chia, and H.C. Toh, Immunotherapy for nasopharyngeal cancer-a review. Chin Clin Oncol, 2016. 5(2): p. 22.

35. Lutzky, V.P., et al., Cytotoxic T cell adoptive immunotherapy as a treatment for nasopharyngeal carcinoma. Clin Vaccine Immunol, 2014. 21(2): p. 256–9.

36. Hui, E.P., et al., Phase I trial of recombinant modified vaccinia ankara encoding Epstein-Barr viral tumor antigens in nasopharyngeal carcinoma patients. Cancer Res, 2013. 73(6): p. 1676–88.

37. Chia, W.K., et al., A phase II study evaluating the safety and efficacy of an adenovirus-DeltaLMP1-LMP2 transduced dendritic cell vaccine in patients with advanced metastatic nasopharyngeal carcinoma. Ann Oncol, 2012. 23(4): p. 997–1005.

38. Lin, C.L., et al., Immunization with Epstein-Barr Virus (EBV) peptide-pulsed dendritic cells induces functional CD8+ T-cell immunity and may lead to tumor regression in patients with EBV-positive nasopharyngeal carcinoma. Cancer Res, 2002. 62(23): p. 6952–8.

39. Teow, S.Y., H.Y. Yap, and S.C. Peh, Epstein-Barr Virus as a Promising Immunotherapeutic Target for Nasopharyngeal Carcinoma Treatment. J Pathog, 2017. 2017: p. 7349268.

40. Hu, L., et al., Comprehensive profiling of EBV gene expression in nasopharyngeal carcinoma through paired-end transcriptome sequencing. Front Med, 2016. 10(1): p. 61–75.

41. Alnabhan, R., et al., Media evaluation for production and expansion of anti-CD19 chimeric antigen receptor T cells. Cytotherapy, 2018. 20(7): p. 941–951.

42. June, C.H., et al., CAR T cell immunotherapy for human cancer. Science, 2018. 359(6382): p. 1361–1365.

43. Morgan, R.A., et al., Cancer regression and neurological toxicity following anti-MAGE-A3 TCR gene therapy. J Immunother, 2013. 36(2): p. 133–51.

44. Cho, H.I., et al., A novel Epstein-Barr virus-latent membrane protein-1-specific T-cell receptor for TCR gene therapy. Br J Cancer, 2018. 118(4): p. 534–545.

45. Yu, F., et al., Presence of lytic Epstein-Barr virus infection in nasopharyngeal carcinoma. Head Neck, 2018. 40(7): p. 1515–1523.

46. Simoni, Y., et al., Human Innate Lymphoid Cell Subsets Possess Tissue-Type Based Heterogeneity in Phenotype and Frequency. Immunity, 2017. 46(1): p. 148–161.

47. Baitsch, L., et al., Exhaustion of tumor-specific CD8(+) T cells in metastases from melanoma patients. J Clin Invest, 2011. 121(6): p. 2350–60.

48. Wherry, E.J., et al., Molecular signature of CD8+ T cell exhaustion during chronic viral infection. Immunity, 2007. 27(4): p. 670–84.

49. Finck, R., et al., Normalization of mass cytometry data with bead standards. Cytometry A, 2013. 83(5): p. 483–94.

